# White matter regions with low microstructure in young adults are associated with white matter hyperintensities in late life

**DOI:** 10.1101/517763

**Authors:** Patrick J. Lao, Robert S. Vorburger, Atul Narkhede, Yunglin Gazes, Kay C. Igwe, Vanessa A. Guzman, Briana Last, Christian Habeck, Yaakov Stern, Adam M. Brickman

**Affiliations:** Taub Institute for Research on Alzheimer’s Disease and the Aging Brain, College of Physicians and Surgeons, Columbia University, New York, NY 10032, United States; Institute of Applied Simulation, School of Life Sciences and Facility Management, Zurich University of Applied Sciences, Wädenswil, 8820, Switzerland; Department of Neurology, College of Physicians and Surgeons, Columbia University, New York, NY 10032, United States

## Abstract

**Background:** White matter hyperintensities (WMH) are areas of increased signal observed on T2-weighted magnetic resonance imaging (MRI) that reflect macrostructural white matter damage frequently observed in aging. The extent to which diminished microstructure precedes or results from white matter damage is unknown. The aim of this study was to evaluate the hypothesis that white matter areas that show normatively lower microstructure are most susceptible to develop WMH.

**Methods:** Five hundred fifty-seven older adults (age: 73.9±5.7yrs) underwent diffusion weighted imaging (DWI) and T2-weighted magnetic resonance imaging (MRI). Diffusion weighted imaging scans were processed into parametric maps of fractional anisotropy (FA) and T2-weighted MRI scans were segmented into WMH. All images were spatially normalized to standard space. A FA template was created to represent normative values from a separate, independent sample of young, healthy adults (N=49, age: 25.8±2.8yrs) and a WMH frequency template was created from the segmented WMH in the older adults. We compared FA values between areas defined as WMH with those defined as normal appearing white matter (NAWM) in the older participants. White matter hyperintensity frequency was binned (0-5%, 5-10%, 10-15%, 15-20%, >20%) and we determined whether WMH frequency bins were different by normative FA values defined in the younger group.

**Results:** Fractional anisotropy values were lower (p<0.001) in WMH regions compared with NAWM regions in the older sample. Areas with higher WMH frequency in older adults had lower FA values in younger adults (5-10%>10-15%>15-20%; p<0.001).

**Discussion:** Low FA values are observed in frank WMH, but FA is also normatively low in regions with high WMH frequency prior to damage. Regions with normatively lower microstructure are more susceptible to future damage from factors such as chronic hypoperfusion or pathology.

## 1. Introduction

White matter hyperintensities (WMH) are areas of increased signal intensity visualized on T2-weighted magnetic resonance imaging (MRI) sequences due to macrostructural changes that lead to decreased lipid and increased water content. The most widely accepted biological model of WMH etiology is that damage to blood vessels leads to chronic hypoperfusion [1], while other biological explanations, such as Wallerian degeneration suggest that primary damage from pathology leads to axonal death [2, 3]. White matter hyperintensities have been linked to suboptimal cognitive outcomes [4, 5], Alzheimer’s disease (AD) risk [6, 7], emotional and motoric dysfunction [8], and risk of later development of stroke [9, 10]. There has been recent interest in understanding the nature of the regional distribution of WMH [11, 12] and the extent to which WMH reflect restricted, discrete damage or a “tip-of-the-iceberg” phenomenon in which the abnormal signal is a harbinger of widely-distributed white matter microstructural damage [13, 14].

While clinical-pathological correlation studies show that radiologically defined WMH are strongly associated with macrostructural damage in white matter secondary to small vessel ischemic changes [15–17], they also suggest widespread diminished microvascular density among individuals with marked WMH, even in white matter that is free of frank radiological abnormalities [14, 18]. More recently, the microstructure around tissue characterized as WMH has been interrogated with diffusion tensor imaging (DTI) [19] to reveal subtle abnormalities in white matter surrounding WMH [20–22]. Longitudinal analyses show relatively diminished markers of white matter microstructure (e.g., fractional anisotropy (FA), mean diffusivity, signal intensity) in normal appearing white matter (NAWM) that subsequently develop WMH *de novo* [20, 23]. These findings suggest a white matter “penumbra” [21] in which microstructural disruption precedes the macrostructural damage represented by WMH, and also suggest that the underlying tissue damage extends beyond the borders of defined WMH.

Another distinct, but not mutually exclusive interpretation can be applied to the fairly consistent observation that WMH are associated with diminished DTI-derived measures of white matter microstructure. Regional differences in white matter microstructure, established during maturation, could render different white matter areas susceptible to damage in later life [24, 25]. That is, in addition to local microstructural damage precipitating the formation of WMH, it is possible that white matter areas in the brain that show normatively lower microstructure may be most susceptible to developing WMH in later life, which can be investigated with normative modeling [26, 27].

The first aim of this study was to confirm that areas that are defined on T2-weighted fluid attenuated inversion recovery (FLAIR) MRI as WMH have diminished white matter microstructure relative to NAWM. Then, by defining the normative white matter microstructure throughout the brain in young, healthy adults (i.e., well before any WMH are typically formed), the second aim of the study was to examine whether WMH in older adults occur in regions that have normatively lower microstructure in early adulthood. We hypothesized that WMH more likely form in areas with normatively lower microstructure.

## 2. Methods

Participants were drawn from young (N=49, mean age ± standard deviation: 25.8±2.8yrs, 14M/35F) and older (N=557, 73.9±5.7yrs, 257M/300F) samples to define a normative FA template and a WMH frequency template, respectively. The use of a young, healthy sample mimics the principle of normative modeling in which normative metrics are defined in a cohort of interest (e.g., young and healthy) to investigate deviations due to pathological conditions (e.g., aging and ischemic injury) [26, 27]. The sample of young adults was a subset of a larger sample enrolled in an ongoing imaging study called the Reference Ability Neural Network study (RANN [28, 29]). The inclusion criteria included native English speaking, strongly right-handed, and at least a fourth grade reading level. The exclusion criteria included MRI contraindications, hearing/visual impairment, and medical/psychiatric conditions affecting cognition. Participants were included in the current analyses if they had undergone diffusion weighted imaging (DWI) and were between the ages of 21 and 30. The goal was to derive normative white matter microstructure values among young, healthy adults. Diffusion weighted imaging data were acquired on a 3.0T Philips Achieva scanner (b-value = 800s/mm^2^ along 56 gradient directions; field of view: 224×224mm^2^; reconstruction matrix: 112×112; 75 slices; slice thickness: 2mm; echo time (TE): 69ms; repetition time (TR): 7645ms) and processed with the FMRIB Software Library (FSL) [30] to produce parametric FA maps for each subject in native MRI space.

The older sample was drawn from an ongoing community-based study called the Washington Heights-Inwood Columbia Aging Project (WHICAP [31]). Participants from WHICAP were included in the current analyses if they had received DWI or T2-weighted FLAIR imaging. MRI data were acquired on a 3.0T Philips Achieva scanner for DWI (b-value = 800s/mm^2^ along 16 gradient directions; field of view: 224×224mm^2^; reconstruction matrix: 112×112; 81 slices; slice thickness: 2mm; TE: 69ms; TR: 8185ms), FLAIR (field of view: 240×240mm^2^; reconstruction matrix: 560×560; 300 slices; slice thickness: 0.6mm; effective TE: 338ms; TR: 3051ms; inversion time (TI): 50ms; flip angle: 50°), and T1-weighted structural scans (field of view: 256×256mm^2^; reconstruction matrix: 256×256; 165 slices; slice thickness: 1mm; TE: 3ms; TR: 6.6ms; inversion time (TI): 724ms; flip angle: 8°).

In the older sample, binary masks of WMH were derived as described previously [32]. Briefly, FLAIR images were skull stripped/brain extracted with FSL. On each FLAIR image, voxels were classified as WMH if the value was 1.2 standard deviations or greater above the mean intensity value of white matter, taken from the brain extracted FLAIR after intensity histogram normalization. All remaining voxels in the white matter (i.e. not labeled as WMH), were classified as NAWM. All segmented images were visually inspected and corrected in the case of incorrect WMH labeling.

In order to compute a WMH frequency template, the FLAIR images were spatially normalized to the T2 template in Montreal Neurological Institute (MNI)-defined standardized space [33], and the transformation matrix was applied to the binary WMH mask (MATLAB2017a, SPM12 toolbox; Figure 1). The spatially normalized WMH masks were averaged to create a WMH frequency template (N=557; e.g., 55 out of 557 subjects have a WMH in a given voxel, making the frequency of a WMH in that voxel = 55/557 = 9.9%; Figure 1).

**Figure 1.**
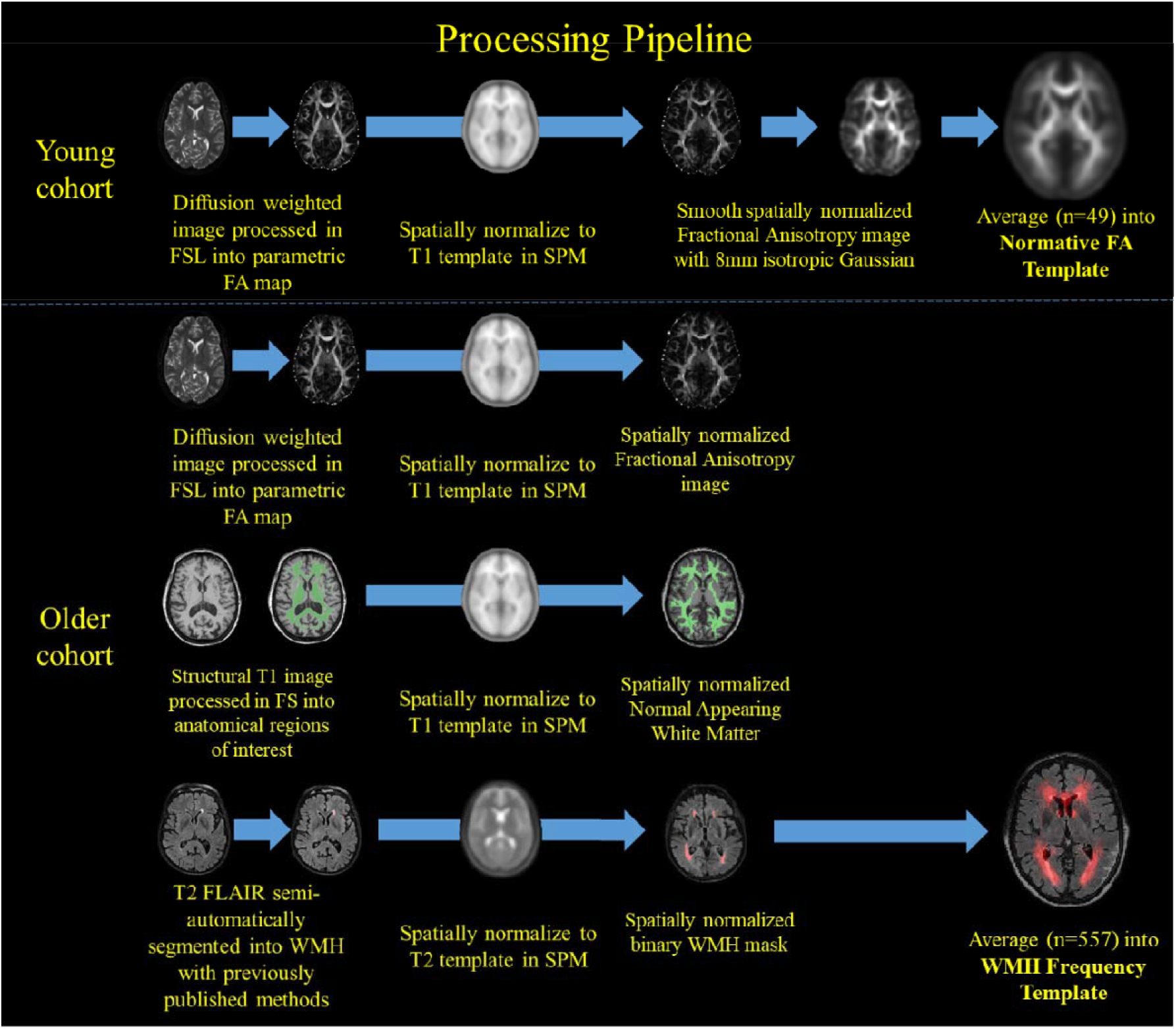
Processing pipeline used to create parametric images and templates for statistical analyses.

In the young sample, parametric FA maps were computed in native MRI space after eddy current and susceptibility corrections using FSL. To produce a normative FA template, the FA maps were spatially normalized to the T1 template in the MNI standard space (MATLAB2017a, SPM12 toolbox; Figure 1). The spatially normalized FA maps were smoothed (8mm isotropic Gaussian) prior to averaging (Figure 1). Parametric FA maps were similarly computed in native MRI space and spatially normalized to MNI space for the older sample (Figure 1).

The comparison of white matter microstructure in WMH and NAWM was performed among the older sample only, where parametric FA maps and binary WMH masks were both available in a given subject. The average FA was computed over all voxels classified as WMH and as NAWM. This approach resulted in two samples for average FA values, one for WMH and one for NAWM, with a common sample size equal to the number of subjects. Distributions were checked for normality, and a paired Student’s t-test was performed to test for a difference in the mean FA between WMH and NAWM.

The WMH frequency template was binned and made into five separate masks representing voxels that were labeled WMH among 0-5% (not including 0%), 5-10%, 10-15%, 15-10%, and >20% (based on the distribution of WMH frequency values which ranged from 0 to 22%). The five binned WMH frequency masks were applied to the normative FA template to derive frequency histograms of FA values in each WMH frequency bin. The number of voxels in each binned WMH frequency mask, as well as the mean and standard deviation for the normative FA template within those WMH frequency masks (e.g., FA values coinciding with WMH frequency values ranging from 0-5%) are reported. The binned WMH frequency masks were applied to the FA maps of the young and older samples, and are presented as the mean and 95% confidence interval. A repeated measures ANOVA with unequal variance (i.e., Welch-Satterthwaite correction) was used to test for differences in FA distribution across WMH frequency bins for the young and older samples, individually. A Gaussian curve was fit to each frequency histogram for visualization purposes only (i.e., mean and standard deviations are from the actual FA values and not from the Gaussian fit of the FA values).

## 3. Results

The normative FA template (Figure 2A) had a range of FA values from 0 to 0.658 with higher FA values overlaying on white matter tracts as expected. The WMH frequency template (Figure 2B) had a range of values from 0 to 22% with a mostly periventricular distribution and with higher WMH probabilities in frontal regions, reflecting the anterior-posterior gradient typically observed in aging.

**Figure 2A.**
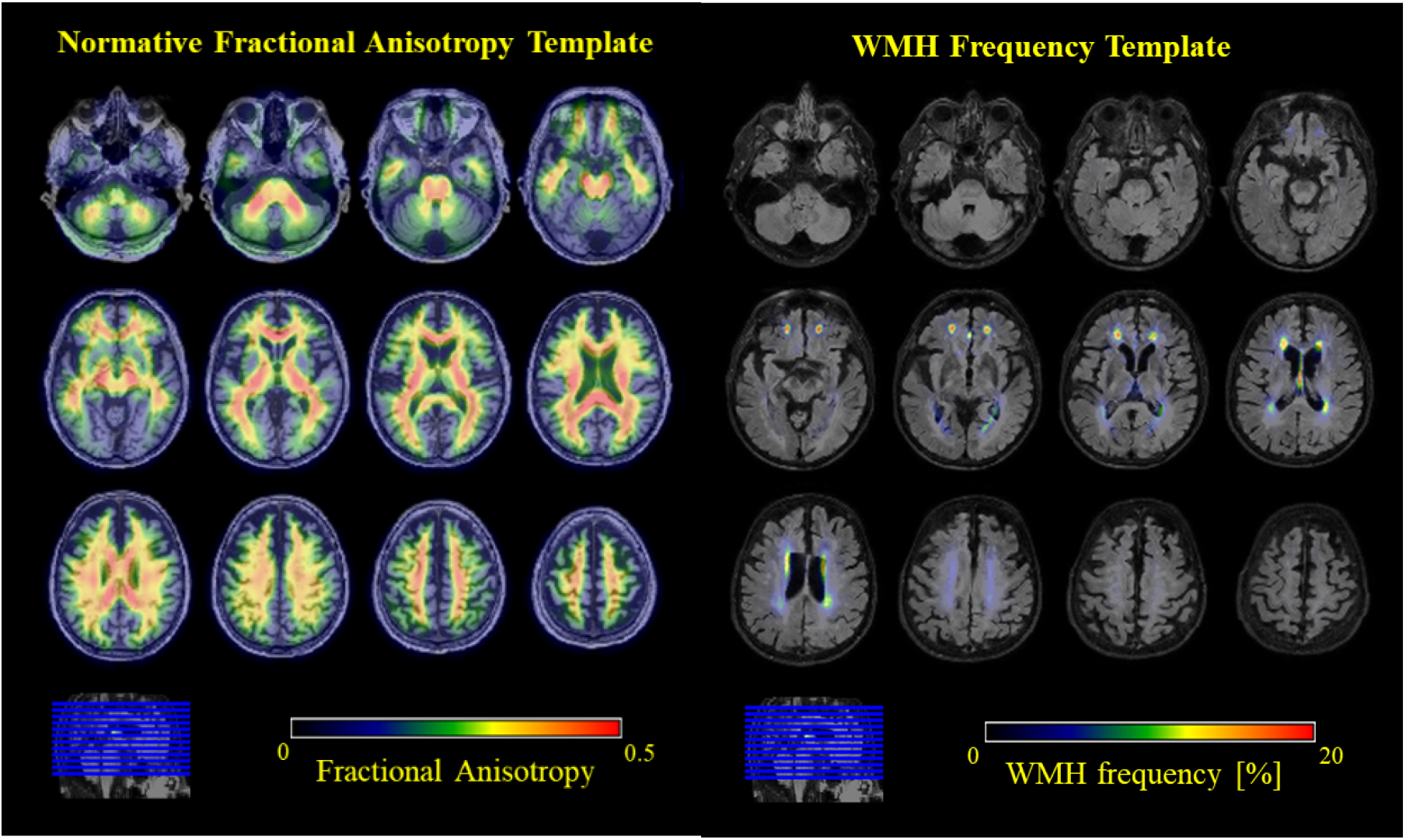
Normative fractional anisotropy template overlaid on a structural image in MNI space. Figure 2B. White matter hyperintensity frequency template overlaid on a structural image in MNI space.

Normal appearing white matter had higher mean FA compared with white matter areas defined as WMH in the older sample (mean difference [95% confidence interval]: 0.017 [0.008, 0.026], p<0.001; Figure 3). When comparing normative FA values as a function of WMH frequency in the older sample, we found that FA values significantly decreased across the 5-10%, 10-15%, and 15-20% WMH frequency bins (F(1.5, 815)=346.1, p<0.001; Table 1). The association did not hold in the 0-5% or >20% WMH frequency bins (Table 1), which were associated with respectively lower and higher normative FA than anticipated. When considering spatial information from the WMH frequency masks (Figure 4), the 0-5% WMH frequency bin contained the highest FA values, but also contained low FA values from crossing white matter tracts or nearby gray matter smoothed into white matter regions. This observation was a consequence of defining this WMH bin by frequencies greater than 0% and lower than 5%, which could include voxels in which only one out of 557 subjects had a WMH. The >20% WMH frequency bin contained only a small number of voxels (n=25) and the FA distribution may not be as reliable compared with those of the other WMH frequency bins (Table 1). While the same limitations exist when looking at the FA distributions in the young sample for WMH frequency bins 0-5% and >20%, it is important to note that there is the same significant decrease in FA values across WMH frequency bins 5-10%, 10-15%, and 15-20% is also observed (F(1.3, 62)=54.1, p<0.001; Table 1) prior to actually developing frank WMH.

**Figure 3.**
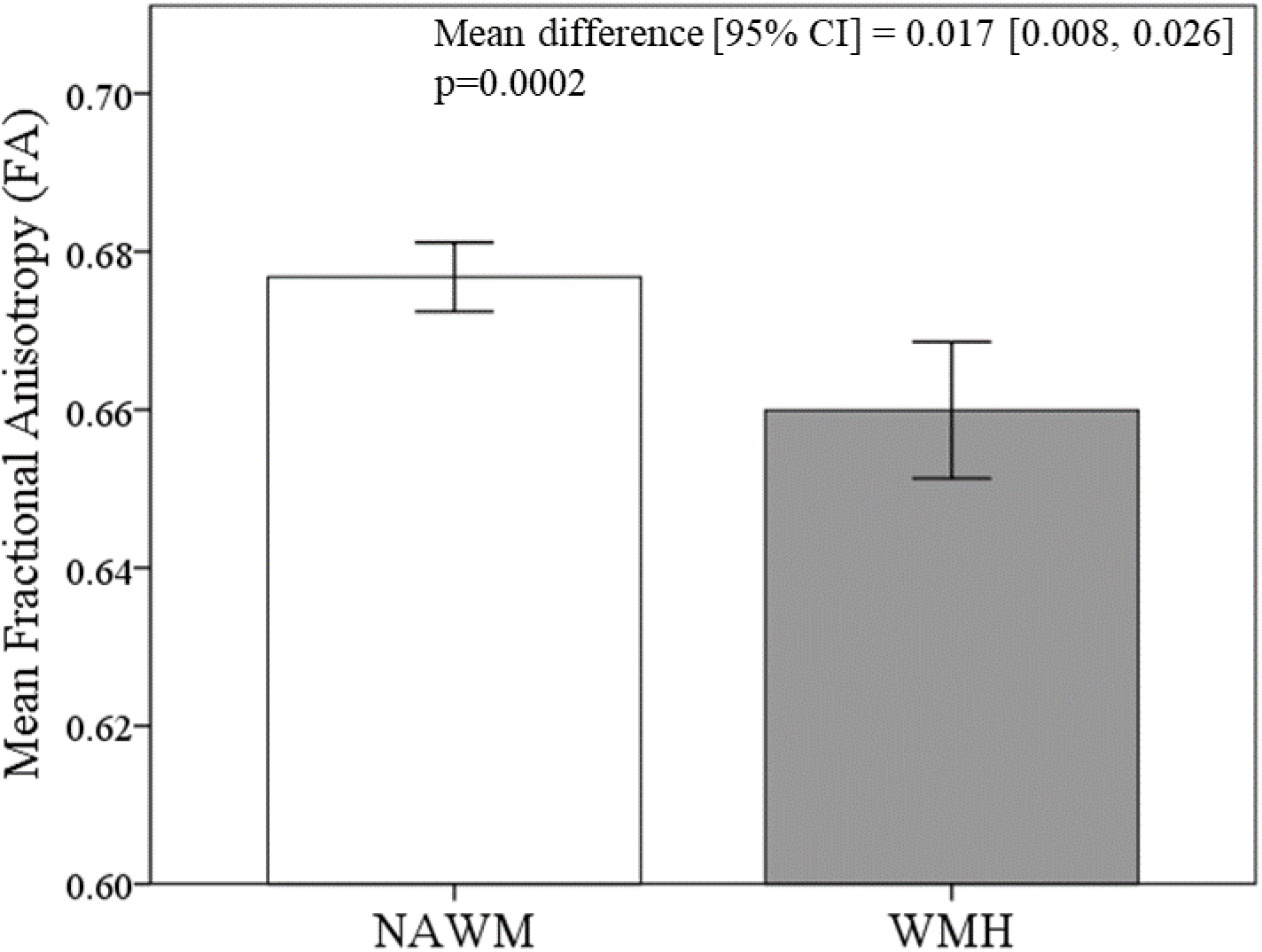
Difference in mean fractional anisotropy value between normal appearing white matter (NAWM) and white matter hyperintensities (WMH) in the older sample.

**Figure 4.**
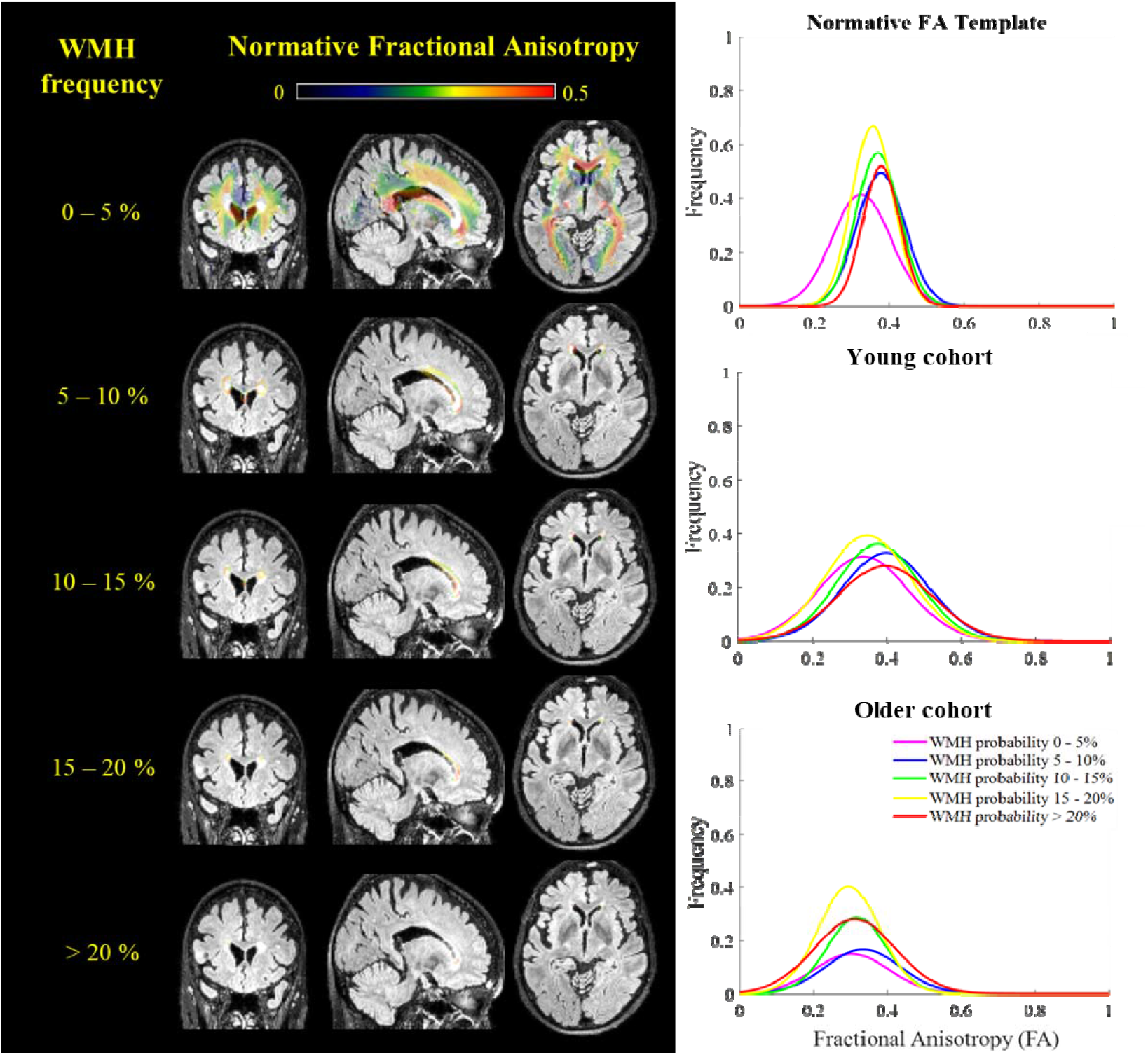
Parametric images of FA values in WMH frequency bins and their corresponding Gaussian fits on frequency histograms of FA values in the normative FA template, the young sample, and the older sample.

**Table 1.** Number of voxels, as well as mean FA and standard deviation for normative FA values or mean FA and 95% confidence interval in the young and older samples, respectively, for each WMH frequency bin.

| **WMH**<br>**Probability Bin** | **Number of**<br>**voxels** | **Normative**<br>**FA** | **Young**<br>**Cohort** | **Elderly**<br>**Cohort** |
| --- | --- | --- | --- | --- |
| 0 - 5% | 53738 | $0.33 \pm 0.08$ | 0.34 [0.33, 0.34] | 0.31 [0.31, 0.32] |
| 5 - 10% | 3812 | $0.38 \pm 0.06$ | 0.40 [0.39, 0.40] | 0.35 [0.35, 0.36] |
| 10 - 15% | 1134 | $0.37 \pm 0.06$ | 0.38 [0.37, 0.38] | 0.33 [0.33, 0.33] |
| 15 - 20% | 316 | $0.36 \pm 0.06$ | 0.35 [0.34, 0.36] | 0.29 [0.29, 0.30] |
| > 20% | 25 | $0.38 \pm 0.05$ | 0.39 [0.38, 0.41] | 0.31 [0.31, 0.32] |

## 4. Discussion

We confirmed that areas defined radiographically as WMH have reduced microstructure, indexed by lower FA values, compared with NAWM in older adults. We further demonstrated that white matter areas with normatively lower microstructure in young, healthy adults coincide with regions that are more likely to appear as WMH in older adults. This study used stringent inclusion/exclusion criteria to recruit a normative cohort of young, healthy adults [26–28] to join previous efforts that have exploited the complementary information provided by DTI measures and WMH [13, 23] in order to understand the relationship between microstructural and macrostructural white matter abnormalities in aging [13, 14, 34].

Among older adults, WMH had decreased FA compared to NAWM. These within-subject, cross-sectional observations could indicate two possibilities, which are not necessarily mutually exclusive. First, macrostructural tissue damage from ischemic injury as measured by WMH [4, 35, 36] results in microstructural white matter abnormalities that are captured by lower FA values on DTI. Second, areas with low white matter microstructure are more vulnerable to ischemic injury in later life. To test the latter possibility, we derived regional, normative microstructural profiles in young, healthy adults and showed that regions with normatively low FA are associated with higher WMH frequency in older adults. The implication of our findings is that regional developmental differences in white matter (e.g., amount of myelin, packing density, size of axons, etc.) could render some areas more vulnerable to ischemic injury in later life. The dependence of late life events on developmental differences is supported by the retrogenesis hypothesis, which postulates that the sequence of degenerative events in the aging brain follows the reverse order of that in the developing brain [24], and that protracted development is driven by myelin producing oligodendrocytes [25]. It might be possible that areas of low white matter microstructure have fewer oligodendrocytes, making those regions more susceptible to macrostructural damage due to chronic hypoperfusion.

Our observation that areas defined as WMH have diminished microstructure compared to NAWM agree with previous reports [37–39]. These findings are also consistent with previous results that observed a “penumbra” in white matter areas within close proximity to WMH that was most vulnerable to later development of WMH [20–22]. Interestingly, those studies suggest that WMH represent a “tip of the iceberg” phenomenon in which white matter microstructure and cerebral blood flow abnormalities extend beyond the identified borders of WMH (2-9mm and 13-14mm, respectively)[14] and portend the later development of frank WMH. Our findings complement these observations by demonstrating that there is lower white matter microstructure potentially decades prior to the development of WMH, and suggest that regional differences in white matter maturation lead to differential susceptibility to chronic hypoperfusion and contribute to the development of WMH.

We introduced a somewhat novel approach to investigate the relationship between white matter microstructure measured with DTI and the macrostructural tissue damage observed as WMH on T2-weighted MRI scans. While former studies relied on two group classification analyses (i.e., WMH or NAWM), the present study additionally used WMH frequency, which are distributed on a continuous scale, enabling a more detailed examination of the relationship between normative white microstructure and macrostructural damage. The ANOVA demonstrated that subtle white matter microstructure differences are apparent in young adults within regions expected to develop WMH in old age, in addition to the paired t-test demonstrating that there are white matter microstructure differences in regions with and without frank WMH in older adults. Importantly, these findings suggest that regions with normatively low FA are more susceptible to future WMH. This negative association between WMH frequency and FA values were only apparent in moderate WMH frequency, most likely due to the small number of voxels in the highest WMH frequency bin and the inclusion of regions with crossing fibers in the lowest WMH frequency bin (Figures 2A, 4; e.g., thalamus, superior longitudinal fasciculus, centrum semiovale [40–42]).

There were several limitations in this study. It should be noted that the purpose of the study was not to compare young and older adults but rather to use information from young, healthy adults to define a set of normative values for white matter microstructure and test whether white matter damage in older adults varies as a function of their distribution. With that objective in mind, white matter microstructure differences were directly tested between WMH and NAWM in the older cohort only, since the young cohort do not have WMH. White matter microstructure differences were indirectly tested between regions with differential WMH frequency (defined by the WMH frequency template derived in the older cohort). The lack of direct comparison between young and older samples obviates the need to harmonize MRI acquisition protocols across the two age groups. However, we acknowledge that the different scanning protocols could bias our findings. While a WMH frequency template was created, the current analyses was restricted to binning the continuous values in an ANOVA as opposed to taking full advantage of the continuous WMH frequency measure in a regression model. This was due to individual FA scans from the young cohort and a single WMH frequency template derived from the older cohort. This WMH frequency template could be used as a prior probability template in future WMH segmentation algorithms. Despite these limitations, this study approximates a decades long longitudinal study by investigating the deviations of an older cohort from a young, normative cohort.

The relationship between the distribution of the underlying FA measures and the occurrence of WMH has been shown in a longitudinal study of older adults [14], but our work shows that the relationship can be observed in adults as young as 20-30 years. The normative FA template was created from representative young, healthy adults. The WMH frequency template was created from a large sample of older adults in which WMH are apparent, and yielded frequencies within the range of previously reported prevalence of small vessel cerebrovascular disease (16-46% in elderly) [43]. Both templates provide valuable information that can inform future studies.

## Acknowledgments

Data collection and sharing for this project was supported by the Washington Heights-Inwood Columbia Aging Project (WHICAP, P01AG07232, R01AG037212, RF1AG054023, R56AG034189, R01AG034189, R01AG054520) funded by the National Institute on Aging (NIA). This manuscript has been reviewed by WHICAP investigators for scientific content and consistency of data interpretation with previous WHICAP Study publications. We acknowledge the WHICAP study participants and the WHICAP research and support staff for their contributions to this study. Additional support for this research was funded by NIA grants and R01AG026158.

